# Pervasive correlated evolution in gene expression shapes cell type transcriptomes

**DOI:** 10.1101/070060

**Authors:** Cong Liang, Jacob M. Musser, Alison Cloutier, Richard O. Prum, Günter P. Wagner

**Affiliations:** Yale Systems Biology Institute, 850 West Campus Drive, West Haven, CT-06516.; Department of Ecology and Evolutionary Biology, Yale University, 165 Prospect Street, New Haven, CT-06520.; Interdepartmental Program in Computational Biology and Bioinformatics, Yale University, New Haven, CT-06511.; Department of Obstetrics, Gynecology and Reproductive Sciences, Yale Medical School, 333 Cedar Street, New Haven, CT-06520-8063.; Department of Obstetrics and Gynecology, Wayne State University, 540 E Canfield St, Detroit, MI 48201.; Yale Peabody Museum of Natural History, 170 Whitney Avenue, New Haven, CT-06511.; Integrated Graduate Program in Physical and Engineering Biology, Yale University, New Haven, CT-06511.; Department of Ecology and Evolutionary Biology, University of Toronto, Toronto ON, Canada M5S 3B2.; Developmental Biology Unit, European Molecular Biology Laboratory, Meyerhofstrasse 1, 69117 Heidelberg, Germany

## Abstract

The evolution and diversification of cell types is a key means by which animal complexity evolves. Recently, hierarchical clustering and phylogenetic methods have been applied to RNA-seq data to infer cell type evolutionary history and homology. A major challenge for interpreting this data is that cell type transcriptomes may not evolve independently due to correlated changes in gene expression. This non-independence can arise for several reasons, such as when different tissues share common regulatory sequences for regulating genes expressed in multiple tissues, i.e. pleiotropic effects of mutations. We develop a model to estimate the level of correlated transcriptome evolution (LCE) and apply it to different datasets. The results reveal pervasive correlated transcriptome evolution among different cell and tissue types. In general, tissues related by morphology or developmental lineage exhibit higher LCE than more distantly related tissues. Analyzing new data collected from bird skin appendages suggests that LCE decreases with the phylogenetic age of tissues compared, with recently evolved tissues exhibiting the highest LCE. Furthermore, we show correlated evolution can alter patterns of hierarchical clustering, causing different tissue types from the same species to cluster together. Using a dataset with sufficient taxon sampling, we performed a gene-wise estimation of LCE, identifying genes that most strongly contribute to the correlated evolution signal. Removing genes with high LCE allows for accurate reconstruction of evolutionary relationships among tissue types. Our study provides a statistical method to measure and account for correlated gene expression evolution when interpreting comparative transcriptome data.

## Introduction

Complex organisms, such as birds and mammals, generally have a much higher number of cell types than so-called “simple” organisms, like sponges and *Trichoplax*. For example, there are more than four hundred cell types recognized in humans (Vickaryous and Hall 2006). In contrast, *Trichoplax*, the morphologically simplest free-living animal, has five to six cell types (Grell and Ruthmann 1991; Syed and Schierwater 2002), and in sponges only about ten to eighteen cell types have been recognized (Ereskovsky 2010; Simpson 1984; Valentine, et al. 1994). Viewed through the lens of animal phylogeny, this observation suggests that animal complexity increased, in part, via the evolution of new cell types (Arendt 2008; Arendt, et al. 2016; Wagner 2014). Thus, understanding the evolution of animal complexity requires investigating cell type evolutionary history and homology.

A promising approach for inferring cell type homology and evolutionary history is the use of hierarchical clustering or phylogenetic methods with RNA-seq data to reconstruct cell type trees (Figs. 1A and 1B) (Achim, et al. 2015; Kin, et al. 2015; Musser and Wagner 2015; Pankey, et al. 2014; Pu and Brady 2010; Tschopp, et al. 2014; Wang, et al. 2011). These are analogous to a gene or species trees, but depict relationships among different cell or tissue types. Homologous cell and tissue types are expected to be more closely related to each other than to other tissues (Fig. 1A and 1B). This pattern has been observed in analyses of mature tissues with distinct functions (Brawand, et al. 2011; Merkin, et al. 2012; Sudmant, et al. 2015). However, the opposite pattern has also been documented (Pankey, et al. 2014; Tschopp, et al. 2014), in which distinct cell/tissue types cluster according to species, rather than by homology, even when the tissue types are phylogenetically older than the species (Fig. 1C). Recent analyses of this phenomenon have suggested the clustering pattern is related to both tissue similarity and the divergence times between the species under comparison (Gu 2016; Sudmant, et al. 2015). However, how and why these factors are related to patterns in the reconstructed tree is currently unclear.

**Fig. 1.**
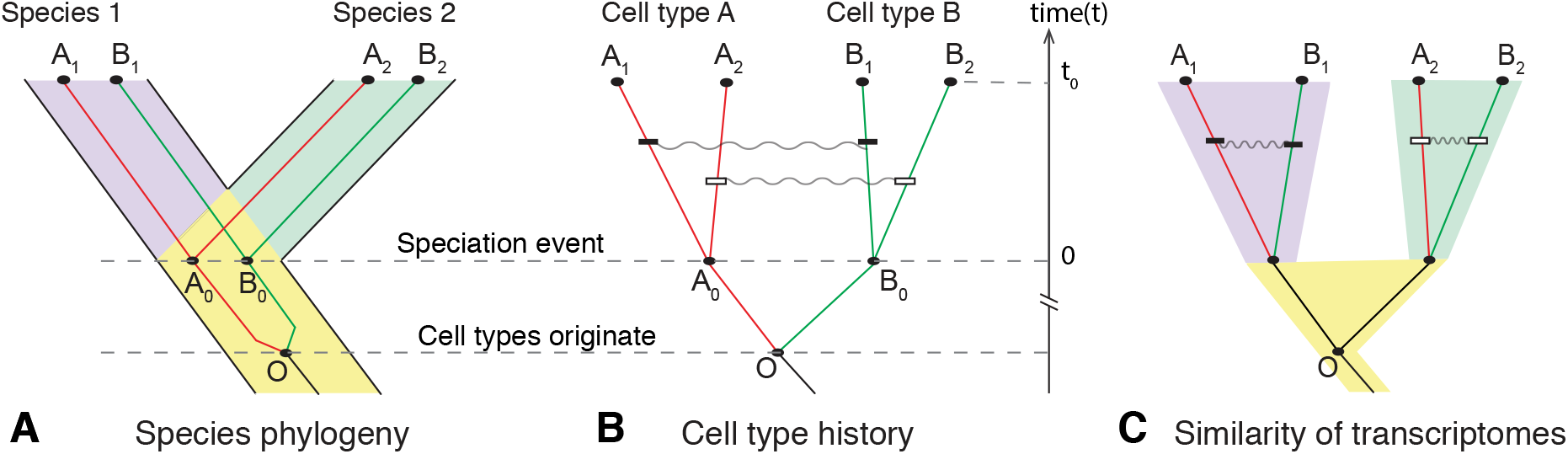
Correlated evolution of cell transcriptomes. **(A)** Cell type history overlaid with species history. Cell types A and B originate from ancestral cell type O prior to split of species 1 and 2. Cell type lineages show homology relationships for the two cell types in the descendant species (subscripts indicate species). **(B)** Cell type history without species phylogeny illustrates cell type homology. Grey squiggly lines indicate correlated changes to the transcriptome. Correlated transcriptome evolution leads to an increase of transcriptome similarities in cell types of the same species relative to their homologous counterpart in other species. **(C)** Correlated evolution causes the accumulation of species-specific gene expression similarities between tissues in the same organism, potentially resulting in different tissues within a species being more similar to each other than to their homologous counterparts in another species.

Phylogenetic inference using tree-building methods assumes that the individuals under comparison (in this case cell types) evolve independently of each other, thus forming distinct historical lineages. Cell and tissue types maintain distinct gene expression programs, and thus may evolve cell and tissue specific gene expression independently. However, many RNA-seq studies find that most genes are not uniquely expressed in a single cell or tissue type. For instance, different neuronal cell types express synaptic genes (Ruvinsky, et al. 2007; Sieburth, et al. 2005; Stefanakis, et al. 2015), different muscle cell types share expression of contractile genes (Brunet, et al. 2016; Steinmetz, et al. 2012), and all tissues employ ‘housekeeping genes’ (Eisenberg and Levanon 2013). Thus, it is likely that some evolutionary changes will alter gene expression across multiple tissues, resulting in correlated patterns of gene expression evolution. This has the potential to influence the relationships inferred in cell type trees by generating similarities that cluster tissues by species, rather than homologous tissue type (Fig. 1B and C) (Musser and Wagner 2015). Correlated evolution of gene expression was recently shown to be important in understanding transcriptome structure using phylogenetic networks (Gu, et al. 2017).

To address the issue discussed above, the level of correlated gene expression evolution (LCE), and its influence on tree reconstruction, needs to be evaluated. Estimation of the correlations among quantitative phenotypic traits along a known phylogeny has been extensively studied both theoretically and with simulations (e.g. Diaz-Uriarte and Garland 1996; Felsenstein 2004; Felsenstein 1988; Felsenstein 1985; Martins and Garland 1991; Pagel 1994; Pagel and Meade 2006). These approaches analyze the evolution of a few traits over a large number of species. In contrast, transcriptomic studies deal with many traits (expression levels of genes or transcripts) over comparatively fewer species, where traditional methods do not directly apply. For this reason, we focus on estimating the average correlation over all genes to obtain an estimate for the overall strength of correlated evolution affecting transcriptomes.

In this study, we propose a statistical method to measure LCE among cell type transcriptomes by incorporating a parameter for correlated evolution in both Brownian and Ornstein-Uhlenbeck (OU) models of gene expression evolution. Using simulations of gene expression evolution, we show that our method accurately estimates the LCE among transcriptomes. Applying this method to published data we find that correlated evolution is pervasive, and varies in accordance with developmental and evolutionary relatedness. We show that LCE decreases with the evolutionary age of the tissues compared, suggesting that the evolutionary independence, or transcriptomic individuation, of tissues increases over time. Further, a theoretical study of our stochastic model of gene expression reveals how LCE and species divergence time influence transcriptome similarities, and thus affect the hierarchical clustering pattern. Using a dataset with sufficient species sampling, we quantify LCE for individual genes. By excluding genes with high LCE, we are able to reconstruct trees that recover expected patterns of cell type homology.

## Materials and Methods

### Publicly available transcriptome data processing

Transcript read counts in Merkin et al. (2012) were downloaded from GSE41637 (http://www.ncbi.nlm.nih.gov/geo/query/acc.cgi?acc=GSE41637). Nine tissues (brain, colon, heart, kidney, liver, lung, muscle, spleen, and testes) in five species (chicken, cow, mouse, rat, and rhesus macaque) are profiled in this work. We followed methods in Merkin et al. (2012) to map each transcript to respective genomes (musmus9, rhemac2, ratnor4, bostau4, galgal3) and extract FPKM values for each gene. We then normalized FPKM to TPM (Transcript per million mapped transcripts) by the sum (Wagner, et al. 2012).

Normalized gene expression RPKM from Brawand et al. (2011) was downloaded from the supplementary tables of the original publication. Six tissues (cerebellum, brain without cerebellum, heart, kidney, liver and testis) in nine mammalian species and chicken are profiled in this work.

Transcript read counts from Tschopp et al. (2014) was downloaded from GSE60373 (http://www.ncbi.nlm.nih.gov/geo/query/acc.cgi?acc=GSE60373). Tail bud, forelimb bud, hindlimb bud, and genital bud in early development stage from mouse and anolis were profiled in this work. Read counts were also normalized to TPM values.

### Skin appendage at placode stage transcriptome data

To collect skin appendage placode transcriptome data we dissected developing skin from embryos of single comb white leghorn chicken and emu. Fertile chicken eggs, obtained from the University of Connecticut Poultry Farm, and emu eggs, obtained from Songline Emu Farm in Gill, MA, were incubated in a standard egg incubator at 37.8 and 35.5 degrees Celsius, respectively. Skin appendages in birds develop at different times across the embryo. We dissected embryonic placode skin from dorsal tract feathers between Hamburger and Hamilton (1951; H&H) stages 33 and 34, asymmetric (scutate) scales from the dorsal hindlimb tarsometatarsus at H&H stage 37, symmetric (reticulate) scales from the hindlimb ventral metatarsal footpad at H&H stage 39, and claws from the tips of the hindlimb digits at H&H stage 36. Dissected skin was treated with 10 mg/mL dispase for approximately 12 hours at 4 degrees Celsius to allow for complete separation of epidermis and dermis. RNA was extracted from epidermal tissue using a Qiagen Rneasy kit. Stand-specific polyadenylated RNA libraries were prepared for sequencing by the Yale Center for Genome Analysis using an in-house protocol. For each individual tissue sample we sequenced approximately 50 million reads (one-quarter of a lane) on an Illumina Hiseq 2000. We sequenced two biological replicates for each tissue, with the exception of emu symmetric scales and claws, for which we were only able to obtain one replicate due to limitations in the number of eggs we could acquire.

Reads were mapped to the respective species genome available at the time: WASHUC2 for chicken (Ensembl release 68 downloaded October 4^th^, 2012) and a preliminary version of the Emu genome generated by the laboratory of Allan Baker at the Royal Ontario Museum and University of Toronto (mapped on December 5^th^, 2012). Reads were mapped with Tophat2 v2.0.6 using the –GTF option. Mapped reads were assigned to genes using the program HTSeq, requiring that reads map to the correct strand, and using the “intersection-nonempty” option for dealing with reads that mapped to more than one feature.

### Estimating the level of correlated transcriptome evolution

For all datasets we calculated TPM (transcripts per million transcripts) as the measure of gene expression level (Wagner, et al. 2012). We also used normalized RPKM from Brawand, et al. (Brawand, et al. 2011) to compare the effect of different quantification methods. All transcriptome data were further processed to facilitate cross-species comparison, with the exception of the normalized RPKM of Brawand dataset which was already normalized between species using a median-scaling procedure on highly conserved genes. For the other datasets we followed the normalization method of Musser and Wagner (2015), using one-to-one orthologs. We then used the square root transformation to correct for the heteroscedasticity of transcriptome data (Musser and Wagner 2015). However, even after the square root transformation we found some very highly expressed genes, such as mitochondrial genes, that still exhibited high variance relative to other genes. These were removed to prevent overestimation of correlated evolution. Finally, we averaged sqrt(TPM) values across replicates to generate a representative transcriptome for each tissue in each species.

We estimated LCE (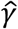) for all combinations of tissue and species in all datasets by calculating the Pearson correlation of their contrast vectors. We tested the significance of correlated evolution by calculating a *p*-value from a null distribution of randomly permutated gene expression contrast vectors. One challenge of estimating the *p*-value is that gene expression evolution between genes is not independent (Ghanbarian and Hurst 2015; Oakley, et al. 2005), for instance because genes can be co-regulated by the same transcription factors (Pavlicev and Widder 2015). To account for this, we subsampled 50 genes for each permutation, which represents a lower bound for the number of independently varying gene clusters identified in single cell transcriptome data (described below). We conducted 1,000 permutations to generate a null distribution of the LCE 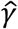. Actual estimates of correlated evolution were considered significantly higher than expected by chance if they fell in the top 5% of the null distribution. We conducted ANOVA tests in R to determine whether correlated evolution values varied depending on time since the last common ancestor of the species used for comparisons. The Welch’s *t*-test was used to determine whether the mean 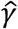 values estimated from the Brawand and Merkin datasets for the same tissue pairs were significantly different.

To estimate the lower bound of independently varying gene clusters, we used the approach of Wagner and colleagues (Wagner, et al. 2008) to estimate an effective number of genes. We randomly sampled different sets of *n* genes. For each set of *n* genes we calculated a correlation matrix using gene expression values from the single-cell transcriptome dataset of Pavlicev, et al. (2017). We then calculated the eigenvalues of the correlation matrix, and the variance of the eigenvalues. According to (Wagner, et al. 2008), the effective number of independent genes equals to *n* minus the variance of eigenvalues of the correlation matrix. We observed that the variance of eigenvalues is always less than 1 when *n* = 50. That is to say, if we randomly draw 50 genes, their gene expression can be considered as independent events.

We estimated the per-gene LCE (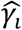) using the correlation between phylogenetic independent contrasts (Felsenstein 1985) for all tissue pairs in the Brawand dataset. We tested its significance using the test statistic 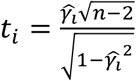. According to the property of the sample correlation coefficient, *t_i_* is distributed as a *t*-distribution with *n* – 2 degrees of freedom under the null hypothesis *γ_i_* = 0. Here, *n* is the number of independent contrasts. We calculated the *p*-value of pergene LCE using the probability of *t* > *t_i_* under the null hypothesis. We calculated the FDR and the fraction of genes under the true null hypothesis (*π*_0_, i.e. the proportion of genes do not have correlated evolution between two tissues) using Bioconductor package ‘qvalue’ (Benjamini-Hochberg procedure). We performed gene ontology (GO) analysis using DAVID bioinformatics tools (v6.8).

## Results

### Stochastic modeling of correlated transcriptome evolution

To model correlated transcriptome evolution, we initially consider two cell types, A and B, in two species, 1 and 2 (Fig. 1 A and B). We assume that the two cell types are more ancient than the last common ancestor of species 1 and 2, which we refer to as species 0. We use vectors *A*_1_, *A*_2_, *B*_1_, *B*_2_ to represent the transcriptome profiles of cell types A and B in species 1 and 2 respectively. Similarly, we use vectors *A*_0_, *B*_0_ to represent the unknown transcriptome profiles in species 0. For each gene *i*, we model the evolution of its expression level in the four cell types as a stochastic process:

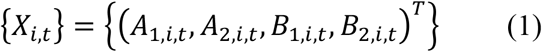

Here, *X_i,t_*’s are multivariate random variables indexed by the continuous time variable *t*, representing the expression level of gene *i* in cell types A and B from two species. We consider the time interval from the speciation event (*t* = 0) to present (*t* = *t*_0_). The initial state of the stochastic process {*X_i,t_*} is *X*_*i*, 0_ = (*A*_0,*i*_, *A*_0, *i*_, *B*_0, *i*_, *B*_0, *i*_)^*T*^. Gene expression values are only measured at the present time *t* = *t*_0_. We studied the stochastic process {*X_i,t_*} with the Brownian model and the Ornstein-Uhlenbeck (OU) model, and developed an estimator of LCE (eq. 6 and 8).

#### Brownian model

We first study the case when {*X_i,t_*} is a Brownian process, a model for neutral transcriptome evolution. The Brownian model is adequate when gene expression changes can be sufficiently explained by random drift (Khaitovich, et al. 2004; Yanai, et al. 2004; Yang, et al. 2017). The accumulated variances in gene expression are proportional to the divergence time in this model. We describe the multivariate Brownian process {*X_i,t_*} with the following stochastic differential equation:

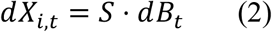

Here, *B_t_* = {*B*_1,*t*_, *B*_2,*t*_, *B*_3,*t*_, *B*_4,*t*_} is a vector of four independent Brownian processes, i.e., they are normally distributed with mean 0 and variance *σ^2^t*. *σ*^2^ is the variance in gene expression that is expected to accumulate per unit time in each cell type. LCE is modeled by parameter *γ* in the correlation matrix *Σ* of this process:

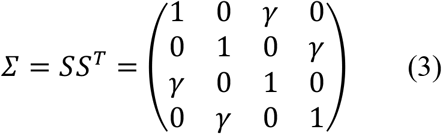

*γ*, ranging in [-1, 1], is the correlation between the gene expression changes in the two cell types: *corr*(*ΔA*_1,*i,t*_,*ΔB*_1,*i,t*_) and *corr*(*ΔA*_2,*i,t*_,*ΔB*_2,*i,t*_). If *γ* > 0, cell types A and B will undergo correlated gene expression evolution. We expect that *γ* > 0 although negative correlations are also mathematically possible. This is because gene expression levels may change toward the same direction if they are co-regulated in two cell types. On the other hand, if *γ* = 0, gene expression changes are uncorrelated in tissues A and B, and no correlated evolution is happening.

We developed a method to estimate LCE parameter *γ* using data from current time, *t* = *t*_0_. For Brownian models, it is known that gene expression vector, *X_i,t_0__* = (*A*_1,*i,t*_0__, *A*_2,*i,t*_0__, *B*_1,*i,t*_0__, *B*_2,*i,t*_0__)^*T*^, is distributed as a multivariate normal random vector with mean 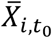 = (*A*_1,*i,t*_0__, *A*_2,*i,t*=0_, *B*_1,*i,t*_0__, *B*_2,*i,t*=0_)^*T*^ = (*A*_0,*i*_, *A*_0,*i*_, *B*_0,*i*_, *B*_0,*i*_)^*T*^, and covariance matrix: *σ*^2^*t*_0_ · *Σ*. Its gene expression contrast vector,

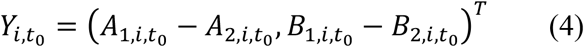

is a linear transformation of *X_i,t_0__*:

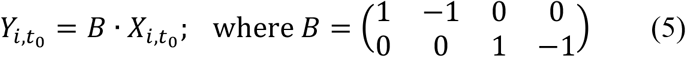

Because a linear transformation of a normally distributed random variable is still normally distributed, *Y_i,t_0__* is distributed as bivariate normal distribution with mean 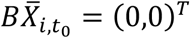 and covariance matrix: 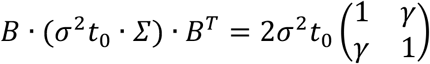, regardless of the initial state of the Brownian process. Thus, we find that *γ* is the Pearson correlation coefficient of the gene expression contrasts between cell types A and B. As a result, we estimate *γ* using the correlation coefficient of the observed transcriptome contrasts:

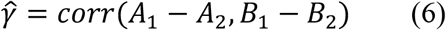

Although we assumed a universal parameter *γ* for all genes in the derivation above, our simulation analysis (see below) showed that when *γ* varies among genes, 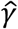 estimated the average LCE among all genes.

#### Ornstein-Uhlenbeck model

We also studied when gene expression evolution is subject to stabilizing selection (Bedford and Hartl 2009; Fay and Wittkopp 2008; Gilad, et al. 2006; Metzger, et al. 2017). In this case, {*X_i,t_*} is a multivariate Ornstein-Uhlenbeck process. This process is described by the following stochastic differential equation:

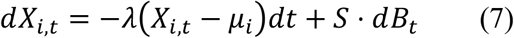

Here, *S* and *B_t_* are defined as the same as in the Brownian model (eq. 3). *λ* represents the strength of stabilizing selection and is always positive. (Brownian model is the limiting case when *λ* = 0.) *μ_i_* = (*μ_A,i_, μ_A,i_, μ_B,i_, μ_B,i_*)^*T*^ is the fitness optimum. For a gene *i*, gene expression optima are the same in homologous tissues, while they may differ between tissues in the same species. Assuming that tissues A and B have evolved long before the speciation event, we study the stationary solution for the OU process.

We took an analogous approach to that we utilized for the Brownian model to estimate LCE parameter *γ* using data from current time. First, as shown in (Meucci 2009), the stationary solution for equation 7 is a multivariate normal distribution with mean *μ_i_* and covariance matrix 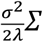. Second, we got the gene expression contrast vector *Y_i,t_0__* through the linear transformation (eq. 4 and 5), and found that *Y_i,t_* is distributed as bivariate normal distribution with mean 0 and covariance matrix 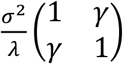, regardless of the evolutionary optimum. Again, we find that *γ* is the Pearson correlation coefficient of the gene expression contrasts between cell type A and B. As a result, we arrived at the same estimator of LCE parameter *γ* as in the Brownian model:

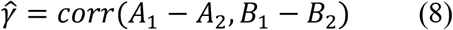

This formula indicates that our estimation of LCE is invariant to some transformations of gene expression measures. For example, if gene expression measure increases by an additive constant *c* in one species, 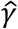 = *corr*(*A*_1_ – *A*_2_ – *c, B*_1_ – *B*_2_ – *c*) remains unchanged.

### Estimation of correlated evolution is accurate in simulated data

We simulated transcriptome evolution data to evaluate our LCE estimator 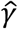 under various conditions. A total of 6119 one-to-one ortholog genes among four mammals and chicken are considered as in Merkin et. al (2012). For each gene, we simulated its gene expression evolutionary trajectory {*X_i,t_*} by iteratively updating *X_i,t_* for a short time period *dt* according to equation 7 (*λ* = 0 for Brownian model; *λ* > 0 for OU model). Negative *X_i,t_* were suppressed to zero in each iteration to avoid negative expression values in the simulation. We used square root transformed gene expression TPMs from Merkin et al. as selective optima *μ_i_* and their initial states *X*_*i*,0_ (see methods). TPMs from two pairs of tissues were used to represent both highly correlated transcriptome profiles (*corr*(lung, spleen) = .74, Fig. 2 blue points) and lowly correlated transcriptome profiles (*corr* (brain, testes) = .24, Fig. 2 orange points). We generated LCE parameter *γ* and stabilizing selection parameter *λ* from various distributions to simulate different evolutionary conditions. *γ* were drawn from two distributions: 1) a beta distribution, and 2) a mixture distribution of beta distribution and zeros. The mixture distribution simulates the scenario that a fraction of the genes experience varied levels of correlated evolution, while the rest of the genes evolve independently. Similarly, *λ* were drawn from three distributions: 1) zeros (Brownian model), 2) a gamma distribution, and 3) a mixture distribution of gamma distribution and zeros. The mixture distribution simulates the scenario that genes are constrained by varied levels of stabilizing selection with a fraction evolves neutrally. We find that our estimation of LCE is robust to varied model parameters. We observed a high correlation (*R*^2^ > .95) between estimated LCE (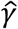) and the mean of true *γ* among genes in all cases (Fig. 2). Meanwhile, as predicted by the theoretical analysis, the correlation in gene expression optima and the initial state of the stochastic process doesn’t influence the estimation of LCE, 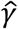 (different colors in Fig. 2). In summary, 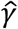 estimated from phylogenetic contrasts accurately estimates the average LCE among all genes under the Brownian and OU process.

**Fig. 2.**
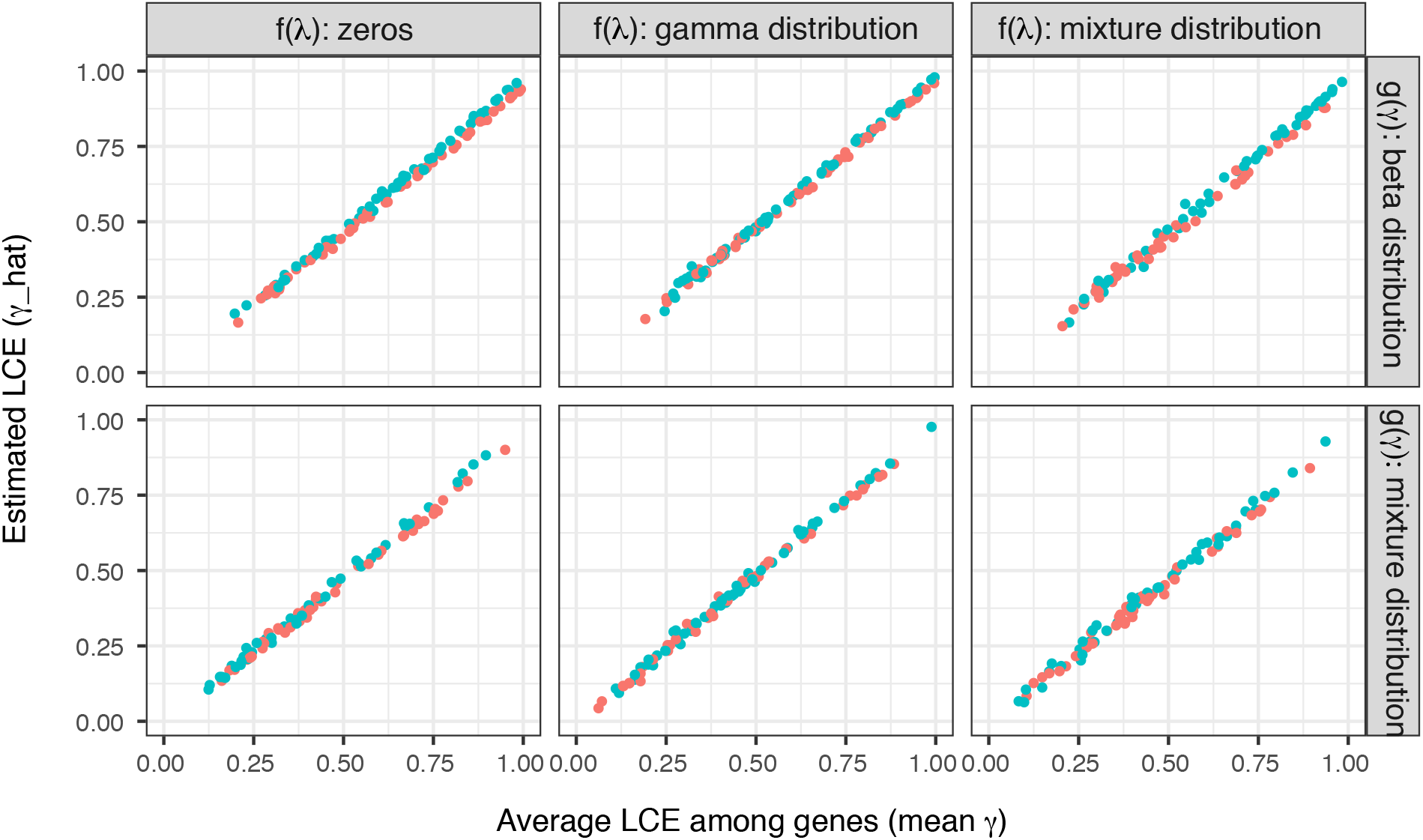
Estimation of LCE in simulated transcriptome data. We simulated 1,000 transcriptome evolutionary trajectories for each condition with varied distributions of stabilizing force *λ*(*λ* = 0; a gamma distribution; a mixture distribution of zeros and gamma distribution) and LCE parameter *γ* (a beta distribution, or a mixture distribution of beta distribution and zeros). In all conditions, we observed a high correlation between the estimated LCE (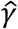) and the mean of true *γ* among genes (*R^2^* > .95). Blue and orange colors are simulations with highly (lung-spleen) and lowly (brain-testes) correlated gene expression optima respectively. We find that estimation of LCE are not influenced by the correlation between gene expression optima or the initial state of the stochastic process. We used square root transformed TPMs from Merkin data as gene expression optima and the initial states in the simulation. A total of 600 data points are randomly sampled to be plotted in this figure.

### Correlated evolution estimation is consistent across datasets

To document LCE among various tissues, we applied our model to three previously published comparative transcriptome datasets: (a) Merkin et al. (2012), containing nine mature tissues from four mammals and chicken (Fig. 3A), (b) Brawand et al. (2011), containing six mature tissues in ten species of mammals and chicken (Fig. 3B), and (c) Tschopp et al. (2014), reporting data on limb, genital- and tail buds at early developmental stages in mouse and *Anolis*. These datasets have three advantages. First, all samples for each data set were generated in the same lab and with the same protocol, reducing batch effects. Second, these datasets include a diverse set of both mature and developing tissues, enabling us to determine the extent to which correlated evolution varies depending on tissue relationships. Third, the Brawand and Merkin datasets sampled five tissues in common (“brain”, heart, liver, kidney, and testes), allowing us to determine whether estimates of LCE are consistent across datasets and species.

**Fig. 3.**
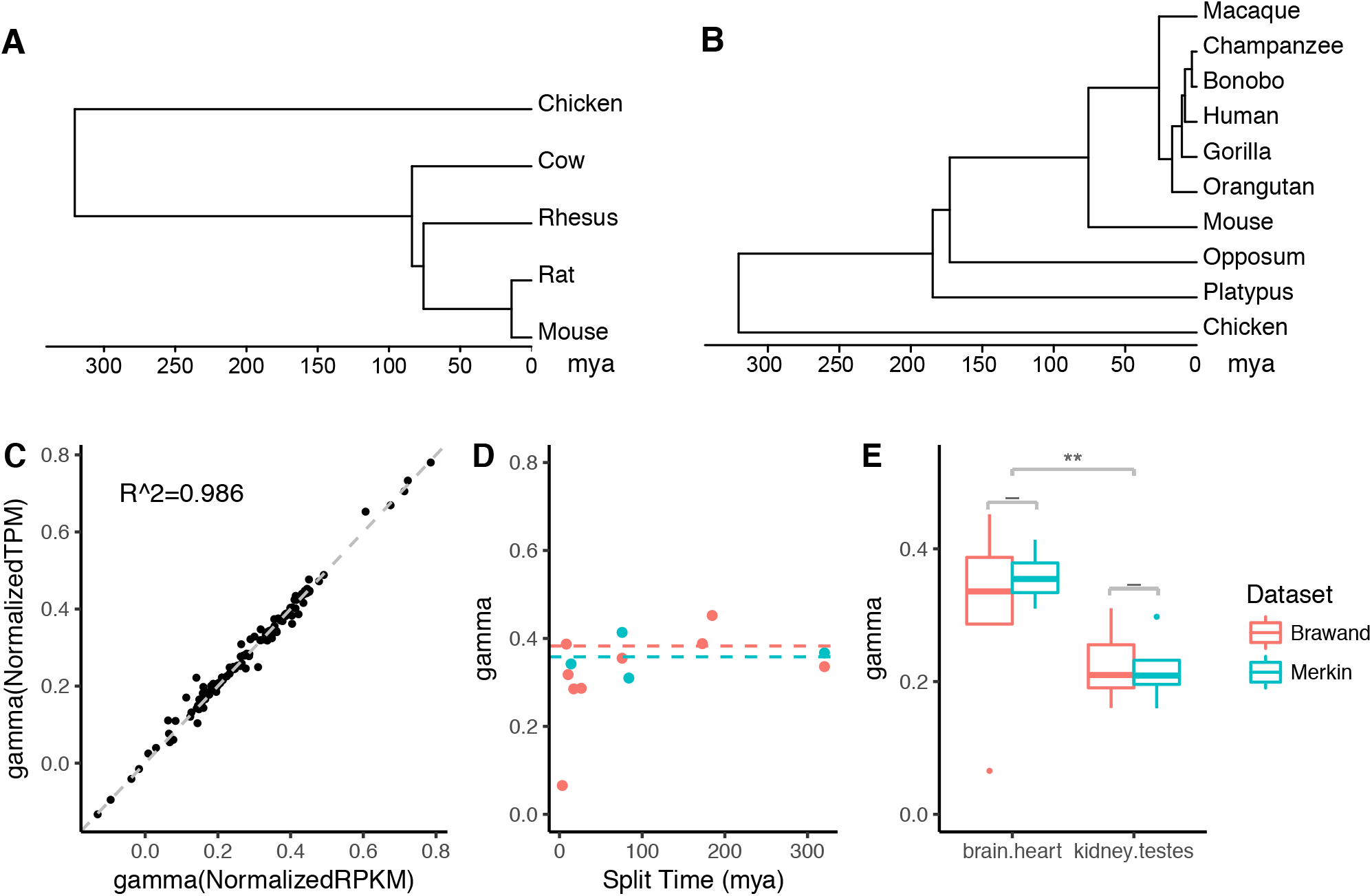
Effect of normalization methods, species divergence time, and dataset on estimates of LCE. **(A)** The time tree for species analyzed in Merkin dataset. **(B)** The time tree for species analyzed in Brawand dataset. All tissues considered here are much older than the species compared in these two datasets. **(C)** The scatterplot of estimated LCE using two normalization methods for Brawand dataset: normalized RPKM according to most consistently expressed genes as in Brawand, et al. (2011), and normalized TPM according to one-to-one orthologs as in Musser, et al. (2015). The normalization methods of gene expression levels do not affect the estimation of LCE (R^2^ = 0.986). **(D)** Estimates of LCE (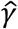) from brain and heart sampled in both Merkin and Brawand datasets using independent contrasts. The horizontal axis is the species divergence time in million years at internal nodes as shown in 3A and 3B. The vertical axis is the estimated LCE, 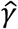. The dashed lines indicate the mean value of 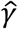 with divergence time > 50mya (blue for Merkin data, orange for Brawand data). The estimated LCE (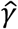) is consistent with species divergence time (ANOVA *p*-value > 0.1). **(E)** Boxplot of estimated LCE for two shared tissue comparisons between Brawand and Merkin dataset (see all shared tissue comparisons in Sup. Fig. 1). Estimates from two datasets do not statistically differ (Welch’s *t*-test, ‘-’: p>0.05, ‘*’: p < 0.05; ‘**’ p<0.01).

We explored the extent to which our estimates of LCE, 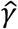, depends on the normalization method of gene expression, the species, and dataset used. To ensure that observations are independent in the following statistical tests, we calculated 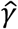 using the correlation between phylogenetic independent contrast vectors (Felsenstein 1985) of gene expression levels in Merkin and Brawand datasets. We found that 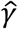 is consistent with different gene expression normalization methods: normalized RPKM as in Brawand, et al. (2011), or normalized TPM as proposed in Musser and Wagner (2015) (Fig. 3C, *R^2^* = 0.986, Sup. Fig. 1A). We found consistent 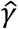 regardless of the divergence time based on estimates by Hedges, et al. (2006) for mature tissues studied in both Brawand and Merkin (Fig. 3D, for all tissue pairs ANOVA *p*-values for dependence of 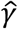 on divergence time > 0.1). However, 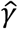 derived from recently diverged species pairs (< 50 million years) tended to be lower, contributing to relatively large standard deviation for some tissue pairs (Table S1-S3). One possible explanation of this finding is that recently diverged species have not accumulated enough gene expression changes to allow for accurate estimation of correlated evolution. Thus, comparisons from recently diverged species may be dominated by observational noise, resulting in underestimation of *γ*. Notably, we found consistent 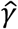 for the same tissue pairs across different datasets (Fig. 3E, Sup. Fig. 1B). The average 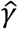 values for the same tissue pair, calculated for Merkin and Brawand datasets separately do not statistically differ (Welch’s *t*-test *p*-values > 0.1), indicating that estimates of the relative LCE are robust to variations in data acquisition. As a result, the fact that our estimation of LCE is consistent with different normalization methods, species, as well as datasets suggests our results reflect biological properties of these tissues, rather than technical artifacts introduced by data acquisition or manipulation.

### Correlated evolution is pervasive and varies in degree across diverse tissues

Our results show that correlated evolution is pervasive and strong, and varies in degree depending on the tissues under comparison (Fig. 4 Table S1 and S2). The highest estimates of LCE were from embryonic appendage buds (Fig. 4; points in red), where 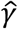 ranged from 0.81 (early hindlimb and tail buds) to 0.89 (early hindlimb and genital buds) consistent with the conclusion of the authors that genital and limb buds are developmentally highly related. Correlated evolution estimates among the differentiated tissues of the Brawand and Merkin datasets were lower. For each pair of tissues in the Brawand and Merkin datasets we calculated 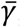, the mean value of 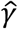 from independent contrasts with split time > 50 million years (Table S1 and S2). Mean estimates ranged from 0.17 (liver and testes) to 0.71 (forebrain and cerebellum), indicating substantial variation in correlated evolution even among differentiated tissues (Fig. 4). Most tissues exhibited at least some degree of correlated evolution, with ∽71% of individual 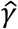 significantly greater than expected by chance (permutation *p*-value < 0.05). Notable exceptions were tissue comparisons with testes, which dominated the lower end of LCE estimates, and could not be statistically distinguished from a model in which gene expression evolved independently between tissues.

**Fig. 4.**
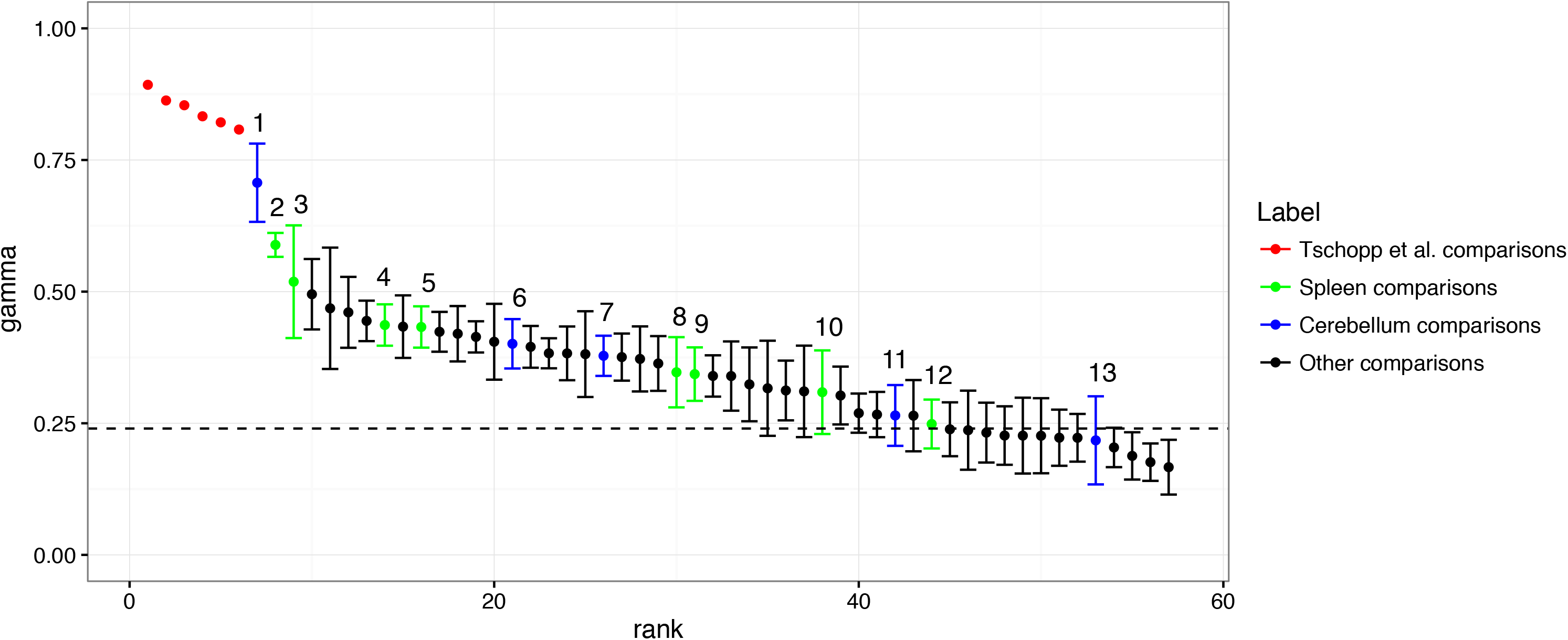
Estimates of LCE in various tissue pairs. LCE estimates for Tschopp, Merkin, and Brawand datasets. Tschopp estimates (*γ*; points in red) are for early forelimb-, hindlimb-, genital, and tail buds from mouse and *Anolis*. Merkin and Brawand estimates (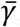) are from 10 different mature tissues in 12 species of mammal and chicken. Estimates of LCE range from 0 to 1, with higher values indicating stronger correlated evolution. Dotted line indicates cutoff for estimates of *γ* significantly greater than expected by chance under a model without correlated evolution. Numbers identify LCE estimates for tissues comparisons with spleen (points in green) and cerebellum (points in blue): 1) cerebellum and forebrain, 2) spleen and lung, 3) spleen and colon, 4) spleen and kidney, 5) spleen and forebrain, 6) cerebellum and heart, 7) cerebellum and kidney, 8) spleen and heart, 9) spleen and liver, 10) spleen and skeletal muscle, 11) cerebellum and liver, 12) spleen and testes, 13) cerebellum and testes.

### Level of correlated evolution differs with phylogenetic age of tissues

The observation that tissues exhibit different LCE indicates that tissues’ evolutionary independence may itself evolve. For instance, do cell and tissue types of recent origin exhibit higher LCE relative to other tissues? To address this question, we collected transcriptomes from developing bird feathers, two different avian scale types, and claws from two avian species, chicken and emu. Feathers are an evolutionary innovation that evolved in the bird stem lineage, replacing scales across much of the body (Prum and Brush 2002). Scales are found in all reptiles, and also on the feet of birds. Bird scales include large, asymmetric “scutate” scales, found on the top of the foot, and small, symmetric “reticulate” scales on the sides and plantar surface. Birds also have claws, which develop from distal toe skin, and develop immediately adjacent to both avian scale types. Claws are shared by reptiles and mammals, indicating that they are phylogenetically older than feathers and scales.

We sampled epidermis during early development of these skin appendages in chicken and emu. LCE estimates (Table 1) ranged from .71 (feathers and claws) to .88 (feathers and scutate scales). Claws, the phylogenetically most ancient skin appendage in this sample, undergo less correlated evolution with respect to the other skin appendages. Feathers, which evolved most recently, are highly correlated with scutate scales (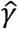 = .88), but exhibit less correlated evolution with reticulate scales (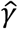 = 79, t-test on bootstrapping *p*=9 10^-4^) or claws (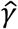 = .72, *t*-test on bootstrapping *p*=3 10^-4^). Notably, feathers and scutate scales undergo more correlated evolution than is found between the two avian scale types (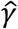 = .86). These results indicate that LCE is not determined by the shared anatomical location of scales and claws on the distal hindlimb. Rather, our data show that feathers evolve in concert with scutate scales. This is consistent with the hypotheses that feathers and scutate scales are serially homologous in early development, and are both derived from ancestral archosaur scales (Di-Poï and Milinkovitch 2016; Harris, et al. 2002; Musser, et al. 2015; Prum and Williamson 2001). Thus, our results suggest that the degree of genetic individuation may increase over evolutionary time, with tissues of more recent origin undergoing relatively higher LCE.

**Table 1.** Estimated LCE among avian skin appendages. Estimates were obtained from transcriptomes of early skin appendage development (placodes) in chicken and emu embryos. For each skin appendage type in each species we averaged gene expression across replicates, and calculated 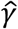 using gene expression contrasts smaller than 10 to avoid the influence of highly expressed structural genes.

|  | **Feather** | **Scutate Scale** | **Reticulate Scale** | **Claw** |
| --- | --- | --- | --- | --- |
| *Feather* | 1 | 0.88 | 0.79 | 0.71 |
| *Scutate Scale* |  | 1 | 0.86 | 0.72 |
| *Reticulate Scale* |  |  | 1 | 0.71 |
| *Claw* |  |  |  | 1 |

### Theoretical analysis of transcriptome clustering patterns

Hierarchical clustering of transcriptomes has been utilized as a discovery tool for cell and tissue type homologies across species (Tschopp, et al. 2014), and for reconstructing the phylogeny of cell type diversification (Arendt 2008; Kin, et al. 2015). However, it is unclear how correlated evolution may influence the result of hierarchical clustering. To explore this, we first conducted a theoretical analysis of how LCE influences the hierarchical clustering pattern of two tissues in two species using our stochastic models of transcriptome evolution. Using Pearson correlation as the similarity measure, samples group by tissue type when the correlation between the same tissue (*corr*(*A*_1_, *A*_2_) and *corr*(*B*_1_, *B*_2_)) is higher than the correlation between tissues from the same species (*corr*(*A*_1_, *B*_1_) and *corr*(*A*_2_, *B*_2_)). Otherwise they group by species. We studied the formula of their transcriptome correlation matrix to predict the hierarchical clustering pattern under various conditions (Fig. 5).

**Fig. 5.**
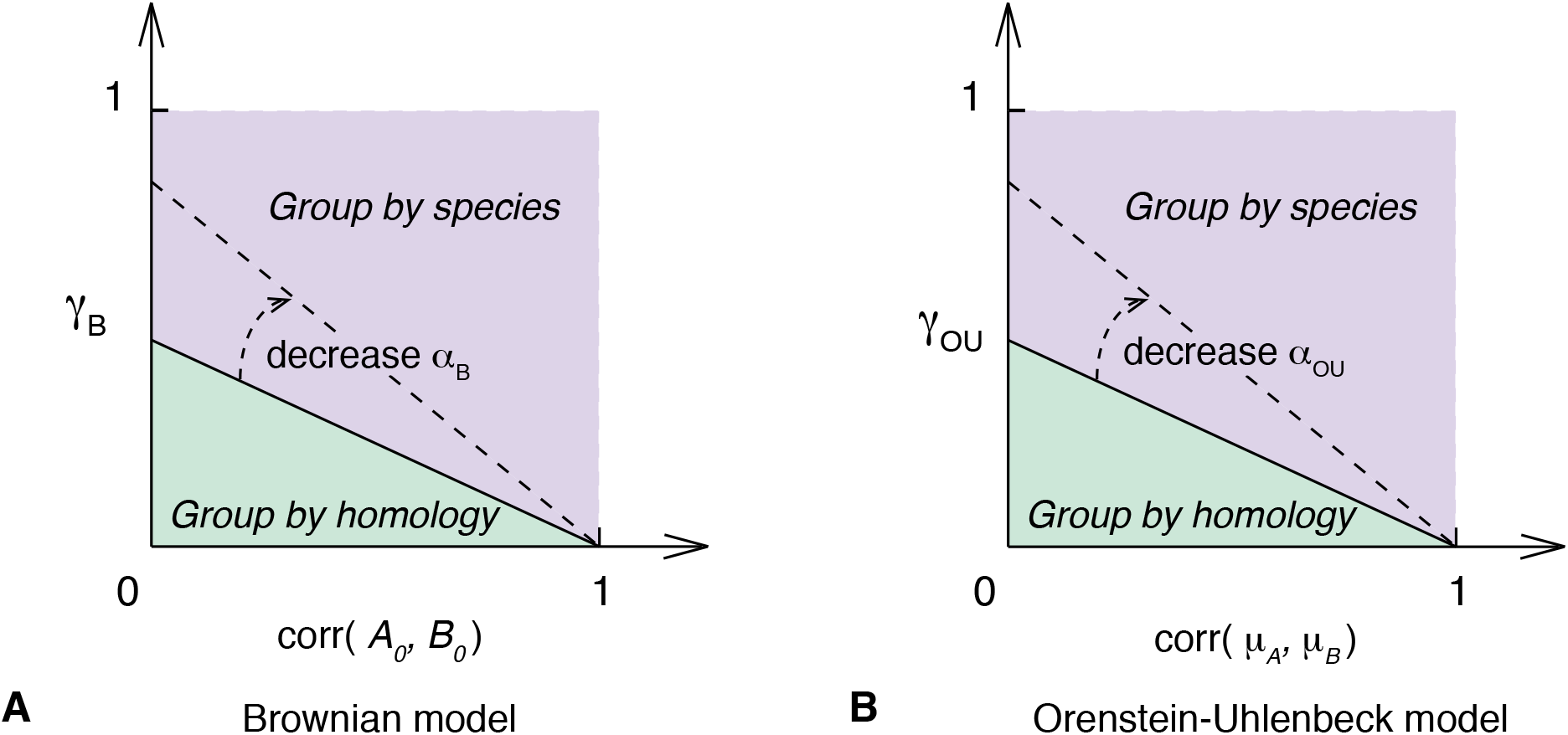
Theoretical hierarchical clustering pattern varies with model parameters. **(A)** Brownian model. **(B)** Orenstein-Uhlenbeck model. The hierarchical clustering pattern of four cell types, A_1_, A_2_, B_1_, and B_2_, is determined by the correlation matrix of their transcriptome profiles (eq. S2). Homology signal (horizontal axis; *corr*(*A*_0_, *B*_0_) or *corr*(*μ_A_, μ_B_*)) and correlated evolution signal (vertical axis; *γ*) shape cell type transcriptome similarities together. The phase transition condition between clustering by homology and clustering by species is a straight line (equations S4 and S5). In both models, clustering by species pattern is observed only with some levels of correlated evolution (*γ* > 0).

Under the Brownian model, the correlation matrix of the four samples is determined by three parameters (Supplementary methods, equations S1-S4): LCE (*γ_B_*), the correlation of gene expression profiles between the ancestral cell types (*r_B_* = *corr*(*A*_0_, *B*_0_)), and the ratio of accumulated random walk variances and the variances in gene expression levels in the ancestral cell types 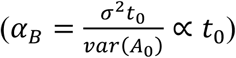. In the parameter space of *r_B_* and *γ_B_*, the phase transition boundary between clustering by species and clustering by tissue is a straight line with positive intersections on both axis (Fig. 5A, Supplementary methods, equations S1-S4). We find that samples cluster by tissue type with small *r_B_* and *γ_B_*. Whereas samples cluster by species when two tissue types are similar to each other ancestrally (large *r_B_*) and have high LCE (large *γ_B_*).

We arrived at a similar conclusion under the OU model. The correlation matrix of four cell types is determined by LCE (*γ_OU_*), the correlation of gene expression optima in two tissues (*r_OU_* = *corr*(*μ*_*A*_, *μ*_*B*_)), and the ratio of stochastic variances and the variances in the expression optima 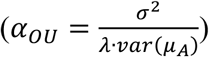 (Supplementary methods, eq. S5). The phase transition boundary between two hierarchical clustering patterns in the parameter space of *r_OU_* and *γ_OU_* is still a straight line (Fig. 5B). We find again that samples cluster by species when two tissue types are similar to each other (large *r_OU_*) and experience high LCE (large *γ_OU_*). In both models, when there is no correlated evolution (*γ* = 0), it is expected that samples group by tissue type.

As a result, the hierarchical clustering pattern is shaped by evolutionary changes as well as historical residuals (Fig. 5). In both models, the correlation between the same tissues from two species reflects, in part, the residual historical signature of homology. However, this historical signal decreases over time due to the variation introduced by the random walk in gene expression evolution. On the other hand, the correlation between tissues from the same species is influenced by LCE, as well as their correlation in the ancestor (Brownian model) or correlation in their fitness optima (OU model). Together, these factors affect the hierarchical clustering pattern of cell type transcriptomes and need to be considered when using comparative transcriptomes in cell type evolutionary study.

### Correlated evolution shapes transcriptome similarities

Our transcriptome evolution model suggests that LCE affects the hierarchical clustering pattern to various extents (Fig. 5). This is consistent with the previous observation that tissue transcriptomes can cluster by species or by tissue homology under different conditions (Gu 2016; Sudmant, et al. 2015). Clustering by species can occur even when the tissues are more ancient than the tissues under comparison, as has been shown in a number of cases (Gu 2016; Lin, et al. 2014; Sudmant, et al. 2015; Tschopp, et al. 2014). Similarly, we observed the species clustering pattern in the re-analysis of three datasets described above, and the pattern persists with different gene expression metrics, including with both normalized TPM and RPKM relative expression values. For example, lung and spleen evolved prior to the common ancestor of birds and mammals, yet we found lung and spleen samples from chicken and mouse clustered by species, rather than tissue identity (Fig. 6A).

**Fig. 6.**
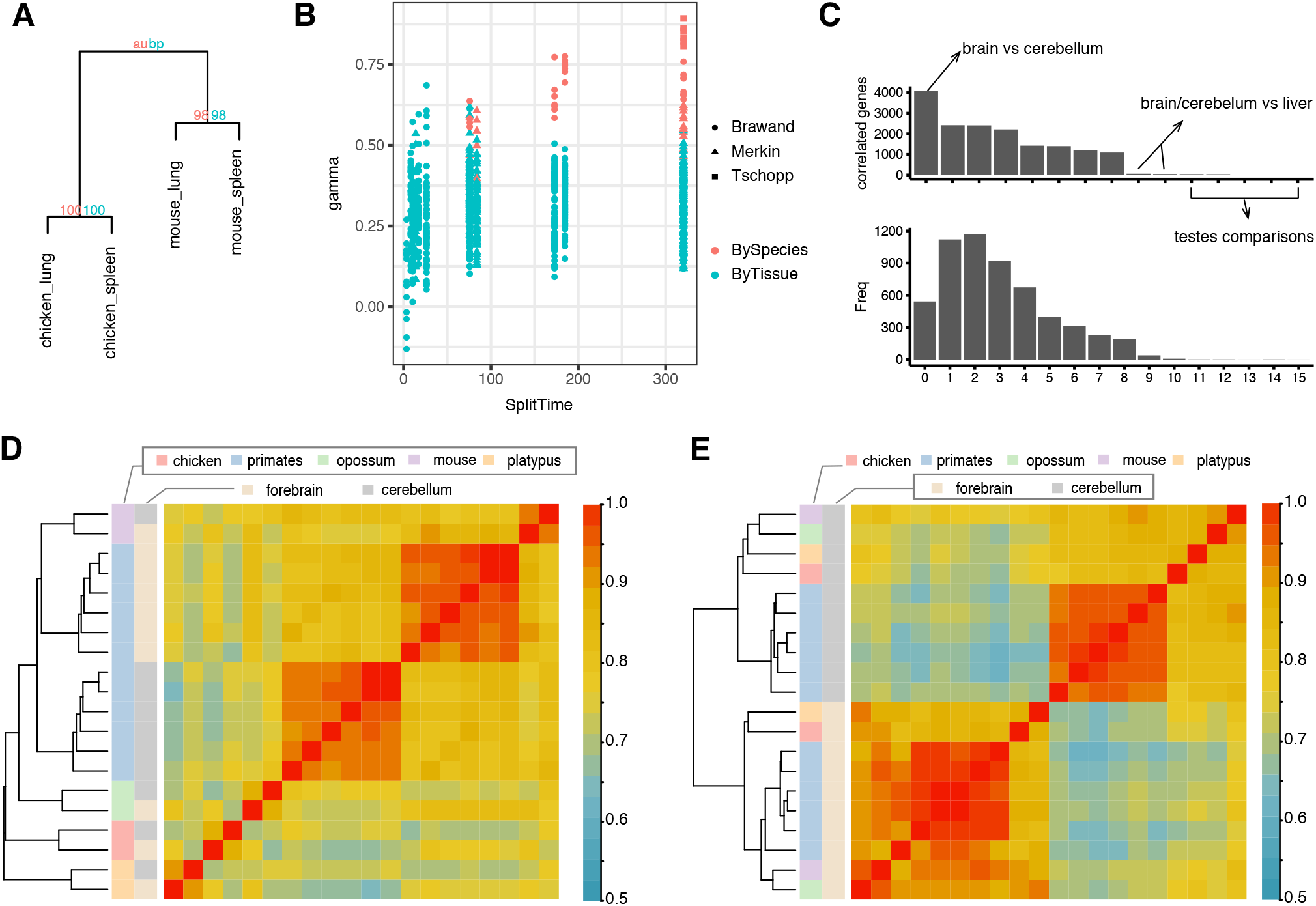
Clustering by homology pattern can be recovered by excluding genes with high LCE. **(A)** Hierarchical clustering of mouse and chicken lung and spleen illustrating examples of tissues clustering by species. Clustering is performed with R package “pvclust”. Numbers at nodes indicate bootstrap support for clusters. **(B)** LCE versus clustering pattern of tetrads with samples from two species and two homologous tissue types. Higher LCE are associated with larger chance of clustering by species. **(C)** Upper: Bar-plot of the number of correlated (high LCE) genes identified in all tissue comparisons in Brawand dataset. The criteria for correlated genes are: *q*-value < 0.05 (BH correction) and 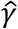 > 0.5. Lower: Histogram of how many times each one-to-one orthologs are identified as correlated genes in all tissue pairs of Brawand dataset. **(D)** Heatmap and hierarchical clustering of forebrain and cerebellum samples from Brawand data using all one-to-one orthologs. Samples cluster by species outside the primates. **(E)** Samples cluster by tissue types when exclude correlated genes in the hierarchical clustering analysis.

To explore the prevalence of clustering by species, and its association with LCE in real data, we determined the clustering pattern for every pair of tissue and species in the Brawand, Merkin, and Tschopp datasets. We found that the majority of mature tissues cluster by tissue type, whereas tissues with high LCE cluster by species (Fig. 6B; symbols in red, Table S4). Further, the more distantly related the species, the more likely tissues clustered by species rather than homology. Tissues collected from species with divergence times less than 50 million years always clustered by tissue homology. However, with more than 50 million years of divergence, nearly all tissue pairs with *γ* > 0.5 clustered by species, rather than homologous tissue. Phylogenetic divergence time also affects clustering because greater divergence time allows for the accumulation of more species-specific similarities via correlated evolution, whereas similarity due to homology remains the same or decays. Thus, in cross species and cross tissue comparisons, transcriptomes reflect both the homology of cell and tissue types, as well as the effect of correlated evolution within each species lineage.

### Clustering by tissue type is recovered by excluding genes with high correlated evolution

The observation above suggest that the effect of correlated evolution needs to be carefully considered when using hierarchical clustering or phylogenetic reconstruction methods with comparative RNA-seq data. To dissect the contribution of each gene to correlated transcriptome evolution, we estimated per-gene LCE (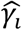) for each one-to-one ortholog *i*, by calculating the correlation between its expression contrasts in two tissues along the phylogenetic tree (Felsenstein 1985). A limitation for estimating per-gene LCE is species number. As a result, we calculated per-gene LCE estimates only for the Brawand dataset, which contained the largest number of species (Fig. 3B). However, we only included gene expression contrasts outside the primate group to avoid biases from dense sampling in one clade and short branch lengths, which could result in inaccurate estimation of variance between species gene expression.

We then tested whether our estimate of per-gene LCE is significant for all tissue pairs (see methods). We estimated the portion of genes that belong to the true null hypothesis (*π*_0_), i.e. do not show correlated evolution, using the *p*-value distribution of all analyzed genes. In Brawand data, *π*_0_ varies from 0.08 (forebrain-cerebellum) to 0.64 (liver-testes) (Table S5), indicating that in this analysis, a large fraction of one-to-one orthologous genes are influenced by correlated evolution. This included 36% of the genes in the liver-testes comparison and 92% in the forebrain-cerebellum comparison. We then identified individual genes with high LCE for each tissue pair using FDR < 0.05 (Benjamini-Hochberg procedure) and with effect size 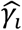 > 0.5. The comparison of forebrain and cerebellum yielded the largest number of highly correlated genes (*n* = 4099), whereas tissue comparisons with testes yielded limited number of highly correlated genes (average *n* = 24; Fig. 6C). The top enriched GO terms in forebrain-cerebellum specific correlated genes (*n* = 631) are related to pan-neuronal functions such as ion transmembrane transport and chemical synaptic transmission (Table S6). We also identified 1190 genes were highly correlated in at least five tissue comparisons. GO analysis of these genes showed enrichment for cell cycle, metabolic and RNA processing related GO terms (Table S7). Finally, we refined the hierarchical clustering analysis of cell type transcriptomes by excluding genes with high LCE. Whereas forebrain and cerebellum samples cluster by species outside the primates using all one-to-one orthologous genes (Fig. 6D), their clustering pattern reveals tissue homology after removing genes with high LCE (Fig. 6E). Hence, identification and elimination of genes with high LCE can be used to recover the phylogenetic history of cell and tissue types.

## Discussion

In this study, we analyzed patterns of transcriptome variation among different cell and tissue types and species using stochastic models of gene expression evolution. We found strong signals of correlated gene expression variation among different cell and tissue types. This pattern of correlated gene expression variation can arise for a number of reasons. The principal ones are: 1) Artifacts in measuring gene expression in different species causing species specific differences in estimated expression level (Gilad 2015; Sudmant, et al. 2015); 2) mechanistic reasons, where either gene regulatory communalities among cell types leads to “concerted” evolution of gene expression due to pleiotropic effects of mutations (Musser and Wagner 2015) or species specific physiological differences affecting gene expression (Lin, et al. 2014) (anonymous reviewer); 3) population genetic causes, where for instance species specific differences in effective population size leads to lineage specific differences in the mutation-selection-drift equilibrium in average gene expression (anonymous reviewer). We address these possible causes of correlated gene expression variation below.

As with any quantitative method, RNA-seq can be subject to artifacts, and some sources of artifacts have been identified (Gilad 2015; Marioni, et al. 2008; Seqc Maqc-Iii Consortium 2014). For instance, batch effects can lead to systematic differences between data sets and thus affect downstream analysis. However, we consider it unlikely that the main result of this study, pervasive and strong correlated variation in gene expression, is caused solely by known artifacts. There are a number of features of our results that are hard to explain as artifacts. One is that the same tissues from different species and data from different laboratories leads to statistically indistinguishable results with respect to the estimated LCE (Figure 3D and 3E). Secondly, artifacts due to different genome quality and transcript annotations should affect estimates of different cell and tissues in the same way, resulting in statistically indistinguishable estimates of LCE between different tissue pairs. In contrast, we find that the LCE varied across different tissue pairs, and is generally higher between tissues with similar cell biology (Fig. 4). For instance, our highest estimates in external datasets were among early developing limb, genital, and tail bud tissue. At this early stage of development, these anlagen largely consist of undifferentiated mesenchymal cells. Moreover, two of the highest measures of correlated evolution between differentiated tissues were for forebrain and cerebellum (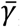 =.71), and heart and skeletal muscle (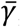 =.47). Both tissue pairs express common sets of pan-neuronal and contractile genes, respectively. Furthermore, we found evidence that developmental origin influences the degree of correlated variation in differentiated tissue. Lung and colon (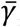 =.50), both endodermal derivatives, were highly correlated relative to other differentiated tissues despite their very distinct functions. These findings along with the fact that our estimation is consistent across varied datasets (Fig. 3) suggest that estimated LCE reflect the biological nature of the tissues under comparison, rather than artifacts of genome quality, transcripts annotation, or batch effects. The overall pattern of gene expression correlations described here would require a quite peculiar set of artifacts to generate the biologically expected differences.

The second possible explanation for the correlated gene expression variation is what has been called “concerted” evolution of gene expression (Musser and Wagner 2015). Concerted gene expression evolution occurs when cell types share parts of their gene regulatory networks. Thus, mutations in certain cis- or trans-regulatory factors can affect gene expression in more than one cell type. This is a special case of pleiotropic effects, a widely recognized pattern for all kinds of characters (Stearns 2010; Wagner and Zhang 2011) and thus plausibly also for gene expression in different cell types. A related explanation is that correlated evolution is expected if species differ in their physiological states in a way that affects gene expression in the two cell types studied. For instance, differences in hormone levels, diet or preferred temperature can lead to species-specific differences in gene expression affecting multiple cells. Correlated evolutionary changes caused by any of these molecular and physiological factors can be called concerted evolution. The term “concerted evolution” originated in the study of nucleotide sequence evolution (Dover 1993; Elder and Turner 1995; Liao 1999), but more generally we use the term here for the “sharing” of mutational effects among different genes, cell types, tissues and organs (Musser and Wagner 2015). This interpretation predicts that tissue specific genes do not contribute to correlated evolution. Consistent with this, we observed significantly decreased LCE when analyzing tissue specific genes only (Sup. Fig. 2, Welch’s *t*-test *p*-value < 10^-6^).

One obvious reason for correlated evolution is the fact that nearly all cell types share certain metabolic and structural “housekeeping” genes, and thus evolutionary changes in their expression would lead to and contribute to correlated gene expression variation. However, our finding that different tissue types from the same set of species (e.g. human and mouse) can have very different LCE (Fig. 3E and 4) suggests that correlated evolution does not solely originate from globally shared “housekeeping” genes. Instead, the correlated evolution signals are heavily influenced by the nature of the tissues and cells compared, suggesting that the structure of the gene regulatory networks active in these cells affects the strength of correlated evolution. For instance, our finding of relatively high LCE between forebrain and cerebellum is consistent with findings in other animals that common transcription factors regulate pan-neuronal gene expression across different neuronal cell types (Ruvinsky, et al. 2007; Stefanakis, et al. 2015). With regards to testes, previous studies have shown testes gene expression has a high degree of tissue-specificity (Lin, et al. 2014; Yue, et al. 2014), is highly divergent among mammals relative to other tissues (Brawand, et al. 2011), and is under directional selection (Khaitovich, et al. 2006; Khaitovich, et al. 2005). Our results suggest that a key factor in explaining testes’ distinct evolutionary history is its ability to evolve gene expression independently, i.e. gene regulatory individuality.

A third possibility is that there are population genetic reasons for correlated gene expression differences (anonymous reviewer, pers. comm.). If mutations, on average, more likely cause a decrease in gene expression (i.e. are deleterious with respect to gene expression) then a decrease in population size will cause an across the board decrease in gene expression level compared to a species that retains a larger population size. That is the case because the mutation-selection-drift equilibrium of gene expression will be lower overall in species with smaller population size than those with higher population size, owing to the fact that selection is less effective in weeding out deleterious mutations in smaller populations. While this is possible, this mode of explanation also predicts equal effects on different cell and tissue types and, as discussed above, the fact that cell and tissue types have different degrees of correlated variation is unlikely to be caused by global factors like population size. Finally, it is a mathematical fact that relative RNA abundance measures, like TPM, the average gene expression levels across genes only depend on the number of transcripts quantifies and thus can not decrease across the board at all (Wagner, et al. 2012).

Overall, it is important to note that the empirical results we report here identify a statistical pattern, correlated gene expression variation between cell and tissue types, which can have a variety of explanations. In this study, we did not specifically investigate the actual cause for these patterns, which needs to be done on a case by case basis. However, our results indicate that this pattern is less likely to be caused by artifacts or some global biological factor. Further empirical investigation is clearly needed for understanding the mechanistic basis for this phenomenon.

### Modeling correlated evolution is essential for reconstructing cell type phylogeny

Our study raises several issues of practical importance for comparative transcriptomics. It is clear that many evolutionary changes in gene expression are not limited to a single cell type, making cross species comparisons difficult. This fact has hampered attempts to use gene expression data to assess the homology of cells and tissues across species, because straight forward clustering of gene expression profiles from different species can lead to obviously wrong results. For example, Tschopp and collaborators (Tschopp, et al. 2014) showed that clustering of transcriptomes is unable to reproduce the fact that limb buds of mice and lizards are homologous. Hence it would be desirable to correct for the effects of correlated variation when attempting to reconstruct the evolutionary history of cell and tissue types from comparative transcriptome data.

Correcting for correlated gene expression evolution is challenging because correlated evolution is occurring at different levels in different tissues. In this study, we outline a potential solution to this problem. We show that correlations of phylogenetic contrasts (Felsenstein 1985), with large enough taxon samples, can identify genes with a high contribution to the correlated evolution signal. Excluding those genes from a hierarchical clustering analysis leads to a result reflecting tissue homology (Fig. 6 D-E). This highlights the importance of sampling more taxa in evolutionary studies of cell or tissue types. Genes that are less vulnerable to correlated evolution in tissues of interest can be identified from a group of species with known tissue homology. These genes allow for the classification of tissues with uncertain homology using unsupervised learning methods, such as PCA and hierarchical clustering (Fig. 6D and E), thus facilitating the study of the origination of novel cell and tissue types. It will be of value to develop analytical tools to systematically correct for the effects of correlated gene expression evolution and thus enable the broad application of the comparative method with respect to cell and tissue transcriptomes.

It will also be useful to extend phylogenetic independent contrasts (PIC) to models other than Brownian motion. Previous simulation results suggest that PIC outperforms other methods under OU model but show decreased accuracy (Diaz-Uriarte and Garland 1996; Martins and Garland 1991). Moreover, there is currently a debate regarding whether Brownian motion is adequate for modelling gene expression evolution (Fay and Wittkopp 2008; Gilad, et al. 2006). Recently diverged taxa, such as among primates, may obey assumptions in the model of Brownian motion (Khaitovich, et al. 2006; Khaitovich, et al. 2004; Yang, et al. 2017). In other cases, transcriptome divergence has been observed to saturate (Bedford and Hartl 2009; Kin, et al. 2016; Metzger, et al. 2017) (Brawand data, Sup. Fig. 3), which may be adequately modeled with an OU model (Supplementary methods, equation S6-7). Thus, A theoretical study of comparative methods under the OU model may advance our understanding of correlated evolution.

## Data access

The raw and processed RNA-Seq data for chicken and emu feather, scales and claw at placode stage are submitted to GEO under accession number GSE89040.

